# A Molecular Rotor-Based Platform for Dissecting Protein Phase Separation and Aggregation

**DOI:** 10.64898/2026.09.22.752683

**Authors:** Jamie Gravell, Nicole Fitikides, Rebecca J. Thrush, Miguel Paez-Perez, Francesco A. Aprile, Ramon Vilar, Marina K. Kuimova

**Affiliations:** Department of Chemistry, Imperial College London, Molecular Sciences Research Hub, White City Campus, 82 Wood Lane, London W12 0BZ, United Kingdom

## Abstract

Biomolecular condensates formed *via* phase separation play critical roles in regulating cellular processes, and their dysregulation is increasingly associated with neurodegeneration. A key challenge in understanding condensate pathology lies in distinguishing dynamic condensates from neurotoxic solid aggregates, the formation of which can, in some systems, occur within existing condensates. Yet current detection techniques often lack the sensitivity and resolution to directly monitor these assemblies and their transitions in dynamic and complex environments. Here, we combined fluorescent probes, based on molecular rotors, with fluorescence lifetime imaging microscopy to quantify changes in protein dynamics during phase separation-associated aggregate formation. Molecular rotors respond to local viscosity changes, which in our study allowed discrimination between fluid condensates and rigid amyloids based on rotors’ fluorescence lifetime. Using α-synuclein, a hallmark protein in Parkinson’s disease, as a model system, we demonstrate real-time visualization of phase separation and condensate maturation within a unified experimental framework. Our method preserves protein integrity and provides quantitative, spatially resolved insights into structural transitions. This technology bridges a critical gap in the study of pathological protein aggregation and offers a powerful platform for mechanistic studies, biomarker discovery, and drug discovery in neurodegenerative disease research.

## INTRODUCTION

Within the complex cellular environment, macromolecules, such as proteins and nucleic acids, can undergo phase separation (PS) to form membraneless compartments termed biomolecular condensates (1). Under normal conditions, condensates compartmentalize biochemical reactions, aiding diverse cellular processes including signal transduction, gene regulation, stress response, and RNA metabolism (2). The functionality of such condensates typically relies on their dynamic and reversible nature, properties maintained by weak, transient multivalent interactions (3-5). In protein condensates these interactions often occur between regions of intrinsic disorder or low complexity. However, dysregulation of condensate nucleation and reversibility has been increasingly linked to aberrant aggregation of proteins linked to several pathologies including neurodegeneration (2, 5, 6).

Neurodegeneration is generally characterized by the accumulation of proteinaceous inclusions in the nervous system (7). These inclusions are composed, in part, of solid protein aggregates, termed amyloids, that exhibit cross-β-sheet structure and are neurotoxic (7). Importantly, amyloid formation can involve intermediate states, including soluble oligomers (8-11) and, in some cases, condensates (12-14). PS can also sequester neurodegeneration-associated proteins, providing an alternate pathway that does not lead to amyloid formation (15, 16). Given the complexity of the PS process, understanding the evolution of the physicochemical properties of condensates over time is essential to elucidate key disease mechanisms, and to develop targeted therapeutic strategies.

While amyloid aggregation has been extensively studied using a range of biochemical and imaging techniques, the characterization and visualization of PS, particularly in relation to disease-associated proteins, remains comparatively less developed. Techniques such as atomic force and electron microscopy, nuclear magnetic resonance, circular dichroism, and fluorescence intensity-based measurements have been instrumental in characterizing the mechanisms of formation and structural properties of amyloid fibrils (17, 18). However, optical microscopy-based methods, alone, provide the combined spatial and temporal resolution required to monitor the physicochemical properties of condensates in heterogeneous mixtures that evolve over time.

Molecular rotors (MRs) are fluorophores that undergo intramolecular rotation or twisting in their excited state. This structural change competes with radiative decay (*i*.*e*., fluorescence). Consequently, low-viscosity environments where molecules rotate freely lead to non-radiative loss of excitation, and thus low fluorescence intensity and short fluorescence lifetimes. In contrast, high-viscosity environments where rotation is hindered result in higher fluorescence intensity and longer fluorescence lifetimes (19). Importantly, lifetime-based microviscosity analysis is well suited to heterogeneous systems, compared to intensity-based techniques, as fluorescence lifetime is concentration independent and so largely unaffected by differences in probe uptake between environments (*e*.*g*., dilute and dense phases or aggregates).

Several MRs have been employed to investigate amyloid aggregation because their fluorescence intensity and lifetime are strongly influenced by the rigid β-sheet architecture of amyloids (20, 21). In fact, one of the most commonly exploited amyloid-binding MRs, Thioflavin-T (ThT), has been used in combination with fluorescence lifetime imaging microscopy (FLIM) to monitor *in vitro* amyloid aggregation kinetics, including the temporal evolution of intermediate oligomer populations (19, 22-24). However, most existing MRs have limited sensitivity and specificity towards assemblies with more dynamic properties, such as liquid-like condensates (25). This limitation underscores a critical gap in our ability to comprehensively characterize the role of PS in disease.

Recently, MR-based FLIM has been utilized to investigate the micropolarity and microviscosity differences of non-amyloidogenic condensates based on multiphasic elastin-like polypeptides, revealing how these properties regulate compartmentalization and maturation (26). Hence, we aimed to design new MRs that are sensitive to, and can discriminate between, the physicochemical properties of both dynamic condensates and solid amyloids important in neurodegenerative disease, overcoming the limitations of existing amyloid-sensitive probes. To do so, we employed a chemically diverse panel of novel MRs with distinct viscosity-sensing characteristics. We assessed their suitability using a FLIM-based strategy that enabled quantitative, spatiotemporally resolved analysis of microviscosity changes during the maturation of condensates into aggregates. As a model system, we applied this approach to monitor PS-associate amyloid formation of the Parkinson’s disease-associated protein α-synuclein (α-syn) (12, 27).

We found that our novel MRs were able to discriminate between α-syn condensates and aggregates, including within their heterogeneous mixtures at intermediate protein aggregation stages. This enables simultaneously real-time imaging of both condensates and aggregates within a single experimental framework, representing a substantial advancement in the study of protein self-assembly. Furthermore, the MRs employed here interact non-covalently with the target assemblies, as opposed to our previously employed strategy for monitoring amyloid β aggregation using MRs covalently bound to the protein (28). This preserves the native behavior of the protein and avoids potential artifacts introduced by direct chemical modification (29). By providing a quantitative strategy to monitor aggregation with high spatial and temporal resolution, our approach opens new avenues for understanding disease mechanisms, identifying biomarkers, and screening for potential new drugs for targeted therapies (11).

## RESULTS AND DISCUSSION

### Selection of condensate- and aggregate-sensitive MRs

This study was motivated by the need for a real-time, high spatiotemporal resolution optical method capable of tracking, within a single experiment, the molecular changes occurring during condensate-associated aggregation. To address this, we developed an approach that enables continuous and direct visualization of these transitions by monitoring the fluorescence intensity and lifetime of MRs during protein self-assembly under PS conditions (**Figure 1a**). To this end, we first analyzed commercially available MRs, ThT and DiSC_2_(3) (**Figure S1**), known to interact with protein aggregates and display fluorescence lifetime sensitivity to different α-syn aggregation intermediates in the absence of PS (22, 23). Additionally, we selected commercial probes ThS (similar to ThT), DilC_2_(3), and sulfo-Cy3-N_3_ (both similar to DiSC_2_(3)) (**Figure S1**).

**Figure 1.**
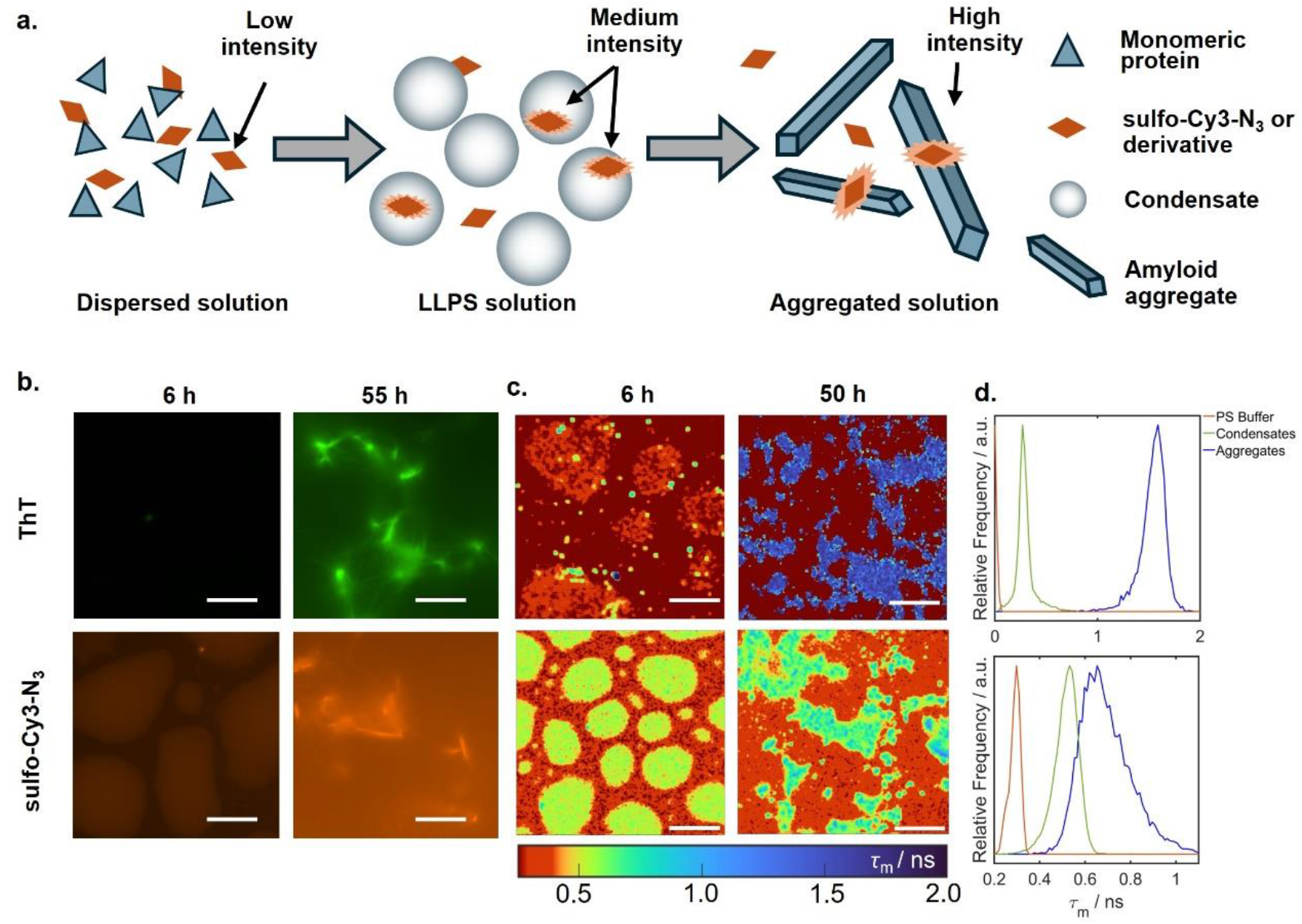
Experimental strategy (a) and characterization (b-d) of condensates and aggregates of α-syn using ThT (top) and sulfo-Cy3-N_3_ (bottom). **a**. Individual α-syn monomers can self-associate to form liquid-like condensates which, under these conditions, are associated with subsequent aggregate formation, which are the hallmark of PD. Both species can be detected using our new family of MRs, while ThT has a limited ability to detect condensates under our experimental conditions. **b**. Fluorescence intensity images of α-syn condensates (∼ 6 h) and aggregates (∼ 55 h) in the presence of ThT or sulfo-Cy3-N_3_. Scale bars represent 25 µm. **c**. FLIM images of α-syn condensates (∼ 6 h) and aggregates (∼ 50 h) in the presence of ThT or sulfo-Cy3-N_3_. The color map indicates the mean fluorescence lifetime measured per pixel. Scale bars represent 100 µm. **d**. Fluorescence lifetime distributions of ThT (top) and sulfo-Cy3-N_3_ (bottom) in PS buffer without protein (orange), or obtained by averaging over all condensates (green) or aggregates (blue) within the FLIM images in **c**.

### Sulfo-Cy3-N_3_ displays high sensitivity for biomolecular condensates

To test the ability of these MRs to probe α-syn condensates, we measured fluorescence intensity and fluorescence lifetime *via* widefield microscopy or FLIM, respectively. This dual approach allowed us to assess MR localization and sensitivity to the physical properties of condensates, *e*.*g*., microviscosity and fluidity. As a model PS system, we incubated α-syn (60 µM) in PS buffer (10% w/v PEG-8000, 0.02% NaN_3_ in PBS, pH 7.4) in the presence of 25 µM poly-L-lysine (PLK, see Methods). These conditions have been shown to consistently induce α-syn complex coacervation and, with time, lead to α-syn aggregation (24, 30). Each coacervate sample was supplemented with either ThT (10 µM), ThS (10 µM), DiSC_2_(3) (3 µM), DilC_2_(3) (1 µM) or sulfo-Cy3-N_3_ (3 µM).

ThT, ThS, DilC_2_(3) and DiSC_2_(3) within α-syn condensates showed minimal fluorescence intensity increase relative to the corresponding free MR in the dilute phase (**Figure 1b** and **Figure S2**). In contrast, sulfo-Cy3-N_3_ exhibited detectable fluorescence intensity increases relative to the dilute phase when inside α-syn condensates or aggregates. This suggests sulfo-Cy3-N_3_ can simultaneously report on a broader range of protein assemblies under the same acquisition settings (**Figure 1b**). Consistent with this finding, FLIM indicated that the fluorescence lifetime of sulfo-Cy3-N_3_ was sensitive to the physical properties of both condensates and aggregates (**Figure 1c,d**). Sulfo-Cy3-N_3_ within condensates exhibited an increased mean lifetime (*τ*_m_(max) ∼ 0.5 ns, see Methods) relative to the dilute phase (*τ*_m_(max) ∼ 0.3 ns). The lifetime distribution of sulfo-Cy3-N_3_ noticeably broadened and shifted towards longer lifetimes (*τ*_m_(max) ∼ 0.6 ns) in the presence of aggregates (**Figure 1d**). In contrast, ThT exhibited a short lifetime within condensates (*τ*_m_(max) ∼ 0.2 ns) which, combined with its very low fluorescence intensity, did not enable reliable condensate detection. However, as expected, the ThT lifetime drastically increased in the presence of endpoint aggregates (*τ*_m_(max)∼ 1.6 ns).

While DilC_2_(3) and DiSC_2_(3) displayed the longest fluorescence lifetimes in the presence of aggregates, both dyes exhibited shorter lifetimes than sulfo-Cy3-N_3_ inside condensates (*τ*_m_(max)∼ 0.3 ns and ∼ 0.4 ns, respectively) (**Figure S3**). However, similar to ThT, the very weak fluorescence intensity of DilC_2_(3) and DiSC_2_(3) within condensates resulted in extended FLIM data acquisition times (∼ 3x longer than for sulfo-Cy3-N_3_), making them unsuitable for studying short timescale changes in condensate properties (*e*.*g*., in response to the addition of small molecules or environmental stimuli). Furthermore, both in the presence and absence of α-syn, DilC_2_(3), DiSC_2_(3), and most significantly ThS, precipitated under our experimental conditions, further highlighting their lack of suitability for our methodology (**Figure S2-4**). Together, these results indicate that of the MRs tested here, sulfo-Cy3-N_3_, which boasts compatibility with high-throughput synthesis, is best suited for detecting and distinguishing protein condensates and solid aggregates.

### Synthesis and characterization of new sulfo-Cy3-N_3_ derivatives

With the aim to further improve the sensitivity of our sulfo-Cy3-N_3_ scaffold (hereafter defined as **1**) for condensates and/or aggregates, we synthesized a panel of seven derivatives *via* click chemistry, specifically Cu(I)-catalyzed azide–alkyne cycloaddition (CuAAC) reactions (**Figure 2a**). Each derivative features a distinct aromatic or heteroaromatic substituent at the triazole ring, selected to modulate the electronic properties, polarity, and hydrogen-bonding potential of the resulting MRs. The newly introduced substituents include pyridine and phenyl groups, as well as five monosubstituted phenyls (with R = -OMe, -NH_2_, -OAc and -CF_3_ in either para or meta positions). Alongside the potential for hydrogen-bonding (as donors or acceptors), the presence of phenyl and pyridine rings may also promote π-π stacking or hydrophobic interactions. We predicted that these features may influence the interaction of our MRs with condensates and aggregates. The new MRs, numbered **9–15** (**Figure 2a**), were prepared using CuAAC reaction conditions optimized to ensure high conversion and purity (above 95%) for all derivatives, as determined by liquid chromatography–mass spectrometry (LCMS) (**Figure S5**).

**Figure 2.**
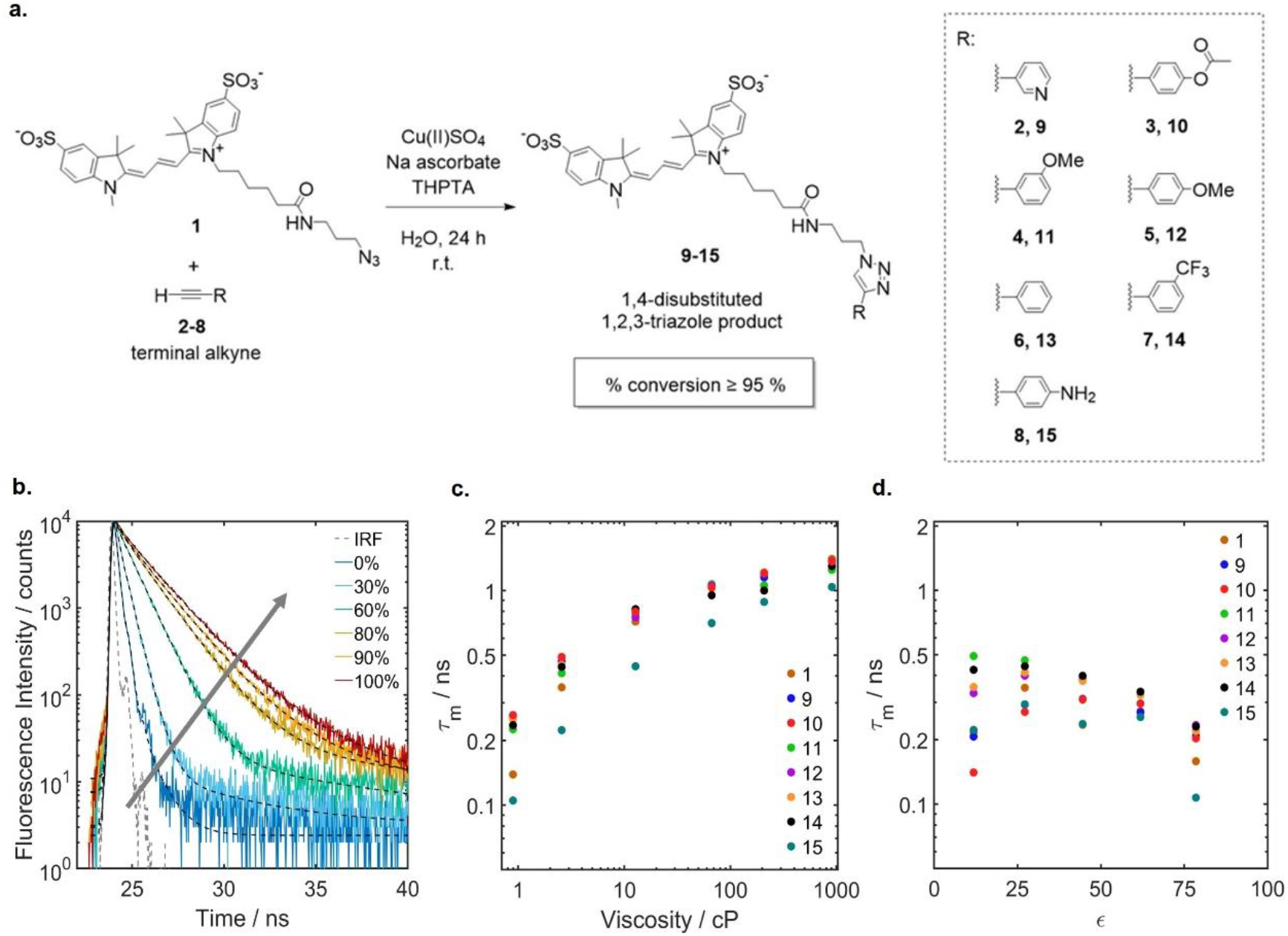
Synthesis and viscosity calibration of the sulfo-Cy3-N_3_ derivatives. **a**. Schematic representation of the synthesis of the sulfo-Cy3-N_3_ derivatives (**9**–**15**) prepared via CuAAC from **1. b**. Time-resolved fluorescence decays of **1** obtained by TCSPC, using 467 nm excitation and 550-560 nm detection in 0-100% glycerol:water mixtures. The effect of increasing viscosity is indicated by a gray arrow. The fits are shown by solid black lines and the IRF is shown for reference. A comparison between the decays of **1** and **9–15** can be found in **Figure S7. c**. Mean fluorescence lifetimes of **1, 9–15** under conditions of increasing viscosity (0-100% glycerol), obtained from tri-exponential fitting of the TCSPC data for **1** (**b**) and **9–15** (**Figure S7**). **d**. Mean fluorescence lifetimes of **1** and **9–15** under conditions of decreasing polarity (0-80% 1,4-dioxane, in a 1,4-dioxane/water mixture), obtained from tri-exponential fitting of the TCSPC data for **1** and **9–15** (**Figure S9**).

We first measured the absorbance and emission spectra of **9–15** and found no notable deviations compared to **1** (**Figure S6**). We then examined the photophysical behavior of **9–15** by using time-correlated single photon counting (TCSPC) to measure their time-resolved fluorescence decay profiles under different conditions. Viscosity effects were assessed using either glycerol/water mixtures of varying ratios (**Figure 2b** and **Figure S7**) or temperature-dependent measurements in 90% or 100% (v/v) glycerol:water mixtures (**Figure S8**). Increasing glycerol content, *i*.*e*., viscosity, increased the fluorescence lifetime of **1** and all derivatives, with each exhibiting sensitivity across a broad viscosity range of ∼ 1 to 905 cP (**Figure 2c** and **Table S1**) (31). All derivatives, excluding **15**, displayed increased fluorescence lifetimes relative to **1** at a given viscosity. **9–15** also exhibited a small temperature sensitivity, showing reduced lifetimes for each viscosity at higher temperature (**Figure S8**). However, this temperature dependence is insignificant compared to the differences in lifetime observed for **1** (**Figure 1c,d**) and **9–15** (*vide infra*) between free in solution and α-syn condensates/aggregates states, for which the samples were incubated at a fixed temperature of 37 °C.

Polarity effects were assessed using 1,4-dioxane/water mixtures, where an increased ratio of 1,4-dioxane:water decreases polarity (quantified by the dielectric constant, *ϵ*). We found that **9–15** also displayed a progressive increase in fluorescence lifetime with decreasing polarity, although all derivatives showed a smaller polarity dependence compared to **1** (**Figure 2d** and **Figure S9**). In fact, the fluorescence lifetime changes over the polarity range studied here were much smaller than those observed when changing viscosity (**Figure S7**), or α-syn amyloid state (*vide infra*). These data indicate that **1** and **9–15** are well suited for detecting changes in viscosity within the condensate environment.

### Functionalization improves sulfo-Cy3 condensate and aggregate sensitivity

We next used widefield microscopy and FLIM to compare the ability of **9–15** to probe α-syn condensates and aggregates, relative to **1**. Throughout the screening process we re-measured the purity of **1** and **9–15** by LCMS. After ∼ 9 months in storage (–20 °C), the purity of each MR was unchanged except for **15**, which displayed a marked decrease (∼ 25%), suggesting that this derivative is unstable. Therefore, **15** was excluded from all further investigations.

**9–14** clearly stained α-syn condensates and aggregates, both by intensity- and lifetime-based microscopy (**Figure 3**). To quantitatively compare the intensity of **9–14** with **1** we applied Otsu’s thresholding to separate each fluorescence image into foreground pixels (*i*.*e*., condensates or aggregates) and background pixels. We then calculated the average intensity per pixel for each MR within each assembly (**Figure 3d**). Each derivative displayed an increase in intensity, relative to **1**, in the presence of either assembly, with **13** displaying the highest intensity within condensates and **14** displaying the highest intensity within aggregates.

**Figure 3.**
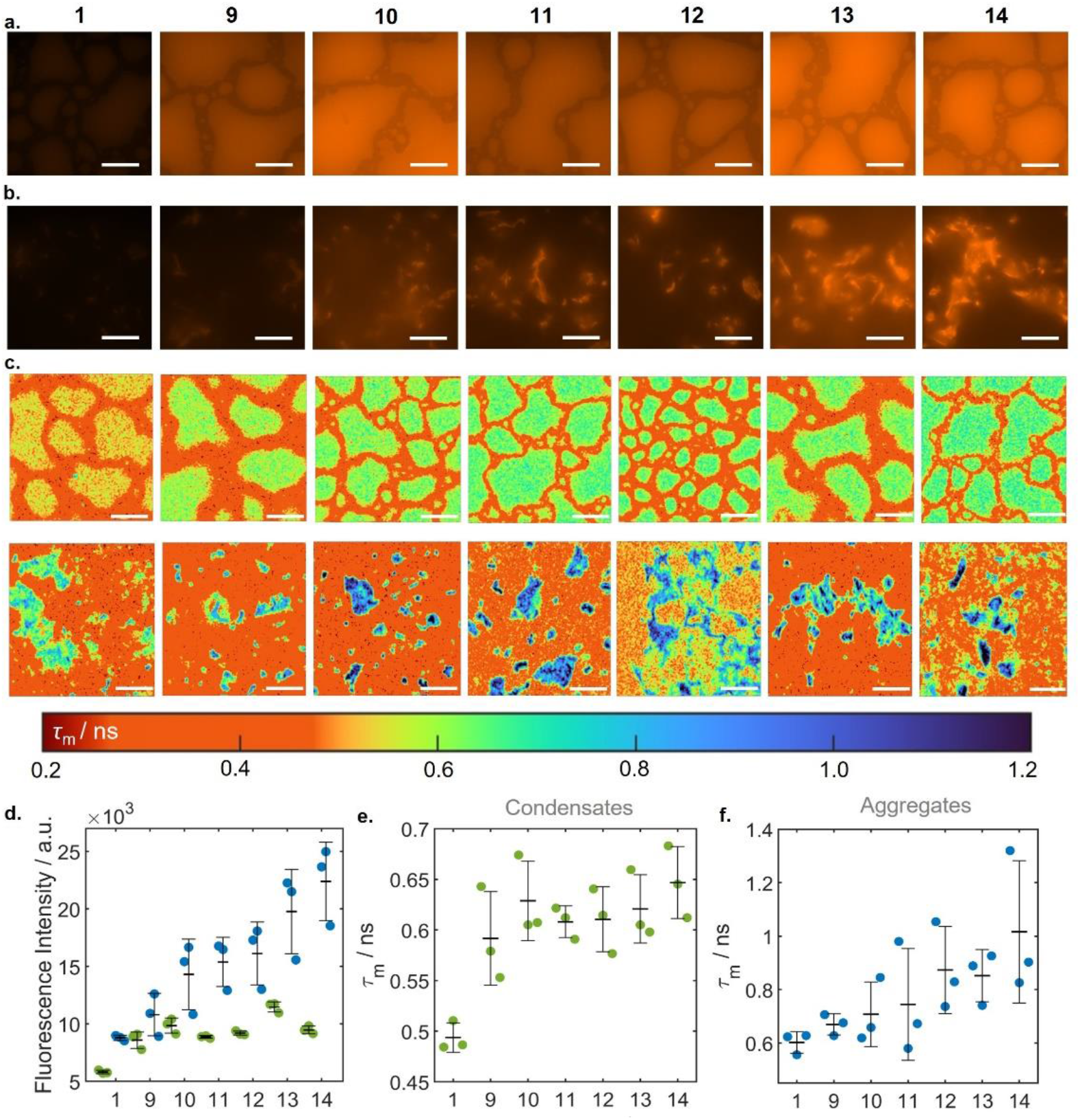
Screening of Sulfo-Cy3-N_3_ derivatives using α-syn PS-associated aggregation. **a/b**. Fluorescence intensity images of α-syn (**a**) condensates (∼ 6 h) or (**b**) aggregates (∼ 60 h) in the presence of **1** or **9–14**. Scale bars represent 50 µm. All condensate images and all aggregate images are processed using the same respective contrast settings. **c**. FLIM images of α-syn condensates (∼ 6 h, top) or aggregates (∼ 60 h, bottom) in the presence of **1** or **9–14**. The false color scale used to represent the mean fluorescence lifetime in all images is also shown. Scale bars represent 100 µm. **d**. Fluorescence intensity quantification of **1** or **9–14** in the presence of α-syn condensates (green) or aggregates (blue). Three biological repeats are shown; each repeat is the mean of three technical replicates. The mean of the biological repeats is shown as a solid black line. **e**/**f**. Quantification of fluorescence lifetime for **1** or **9–14** in the presence of (**e**) condensates or (**f**) aggregates. These values are calculated from the maximum of the lifetime distributions of either assembly in the presence of **1** or **9–14**. Each point represents one biological replicate; the solid black lines indicate the average. Error bars in **d-f** indicate the standard deviation.

To quantitatively compare the lifetime of **1** and **9–14** within condensates and aggregates, we plotted the fluorescence lifetime distribution (**Figure S10**) of each MR in the presence of either assembly, and calculated the corresponding *τ*_m_(max) (**Figure 3e**,**f**). The *τ*_m_(max) of **9–14** increased relative to **1** within both condensates and aggregates, with similar condensate-associated *τ*_m_(max) values for all derivatives and an overall increase in *τ*_m_(max) going from **9** to **14** in the presence of aggregates (**Figure 3e,f**). Furthermore, we observed broader lifetime distributions for all MRs in the presence of aggregates, as compared to condensates, possibly reflecting the presence of multiple, structurally distinct domains within aggregates (**Figure S10**).

The increase in both fluorescence intensity and lifetime for all MRs in the presence of aggregates, relative to condensates, is consistent with the higher viscosity/packing within solid assemblies over more liquid-like condensates, and in non-crowded solution (*i*.*e*. in PS buffer alone). The lifetime of **9–14** also increased, relative to **1**, when MRs are free in solution, consistent with our TCSPC calibration data (**Figure 2c**). The effect of individual modifications to the MRs through derivatization has the most significant impact on the lifetime values obtained in aggregates. Dyes **9–11** offer little advantage, within error, in studying aggregates by lifetime compared to **1**. The lifetime distribution of **14** is the broadest, leading to a large spread in obtained contrast (lifetime) values. Given this, we believe **13** would be the most optimal probe to distinguish between condensates and aggregates, as the lifetime values obtained in aggregates have a narrower spread than **14. 13** also has the highest fluorescence intensity within condensates, allowing shorter FLIM acquisition times and thus can be used to investigate changes in condensate properties on the shortest timescale.

Most of our new derivatives have improved fluorescence intensity and lifetime contrast between condensates and aggregates, enhancing our ability to identify and distinguish these assemblies throughout the PS-associated aggregation pathway. Combined with FLIM, our new MRs provide a simple optical microscopy-based framework to monitor microviscosity changes during condensate maturation with high spatiotemporal resolution.

### 13 can monitor aggregate formation in condensates in a spatiotemporally resolved manner

Having established our approach can distinguish condensates from the dilute phase or aggregates, we then verified that our derivatives could follow aggregate formation within condensates in real time using FLIM. We chose **13** for this purpose as it exhibits the optimal properties for our methodology. α-Syn PS samples were imaged at regular intervals during condensate maturation (**Figure 4a**). We found that initially homogeneous, relatively low viscosity condensates (2h and 4h) begin to mature aggregates (∼ 7-9.5 h) eventually yielding large structures whose lifetimes are consistent with those of endpoint aggregates (**Figure 3c**).

**Figure 4.**
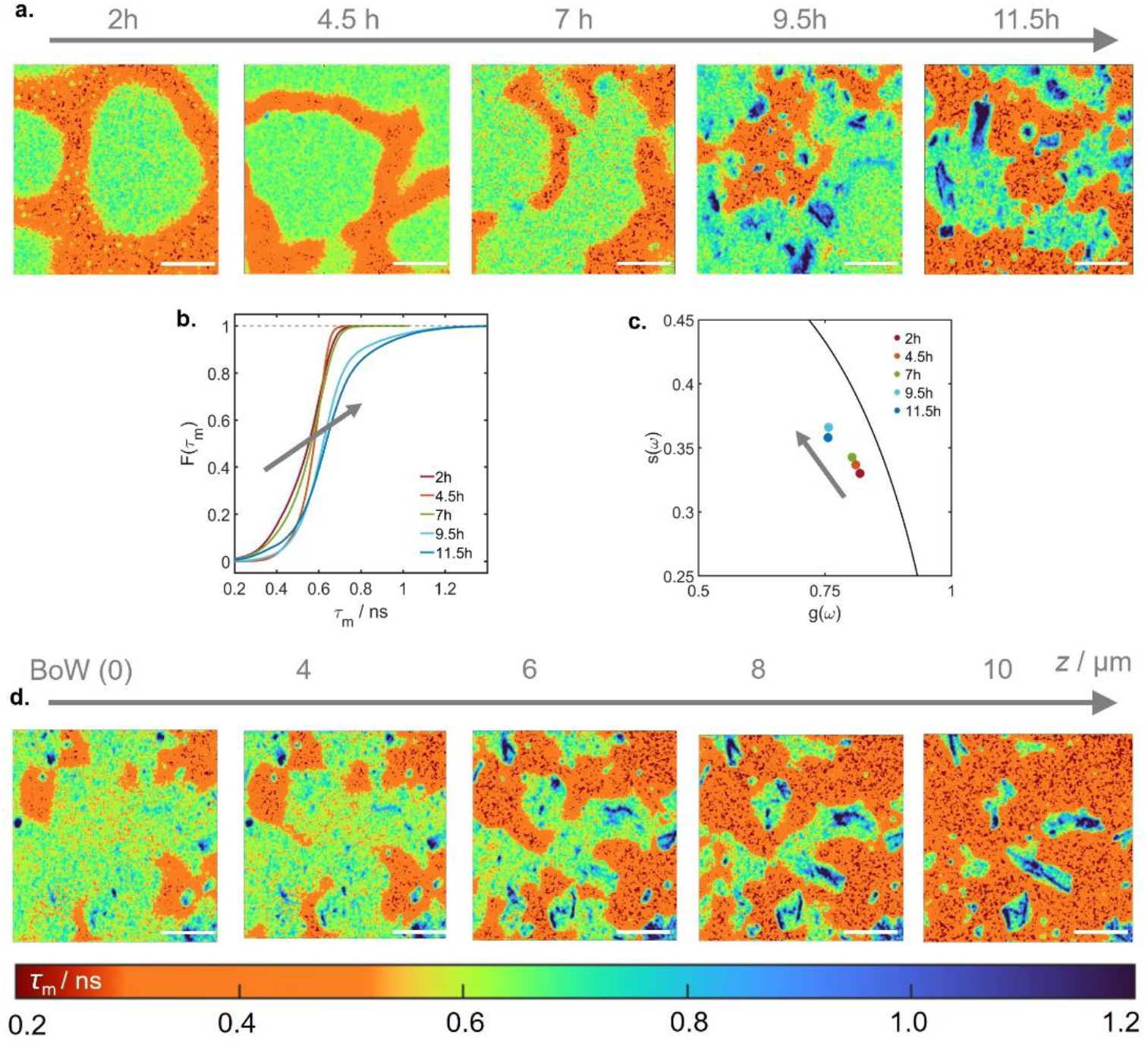
Application of 13 to study PS-associated aggregation in real time. **a**. FLIM images of α-syn condensates in the presence of **13** acquired at 2, 4.5, 7, 9.5, and 11.5 h after induction of PS. **b**. Empirical cumulative distribution functions of the fluorescence lifetime over the segmented condensates from the images given in **a. c**. Phasor plot of the lifetime images presented in **a. d**. FLIM images obtained for α-syn incubated in PS buffer for 11.5 h in the presence of **13**, at increasing distance (left to right) from the silica interface at bottom of well (BoW). The false color scale used to represent the mean lifetime in all images is shown below. All scale bars represent 100 µm.

To quantify aggregation within α-syn condensates, we used the empirical cumulative distribution function (eCDF) of the lifetimes (**Figure 4b**). These functions clearly show a shift to both higher average lifetime and skew of the lifetime distributions towards significantly longer lifetimes upon α-syn aggregate formation. These results were highly reproducible both in terms of kinetics and relative spatial distribution of aggregate within a condensate, occupying primarily the periphery of the droplets which likely provide aggregation sites (32). We could also follow the progression of aggregate formation by employing phasor analysis (**Figure 4c**) - a fitting free, Fourier transform-based approach we have previously employed to monitor amyloid aggregation in the absence of PS. In brief, the real and imaginary components of the Fourier transform of the time-resolved fluorescence decay at each time point during condensate maturation are computed and plotted on the so-called “universal circle” (33). For the FLIM images shown in **Figure 4a**, all these points lie within the universal circle indicative of a multi-component fluorescence decay. Meanwhile, the counterclockwise shift with time indicates an increase in lifetime as PS progresses. It is noteworthy that this analysis method also indicates an increase in condensate lifetime, and therefore viscosity, before large aggregates become visible, indicating that FLIM combined with **13** is well suited to monitor the early stages of protein aggregation under PS conditions.

To our knowledge, this is the first demonstration of a technique capable of selectively monitoring changes to both the condensate and aggregate environments, simultaneously. Relative to other techniques (*e*.*g*. Fluorescence recovery after photobleaching (FRAP)), it also dispenses the need to average over a large region of interest within the condensate, thereby conflating the two environments. Of note, previous FRAP measurements showed an apparent increase in condensate viscosity during aging and aggregate formation, while our measurements show that the MR lifetime within condensates is relatively unaffected in regions without maturing aggregates.(12) We therefore believe that our technique combined with FRAP can offer complementary information on PS-associated aggregation. Our approach also allows details on the aggregation mechanism within condensates to be directly visualized. In this case, we observe preferential aggregate formation near the condensate-dilute solution interface, most clearly seen *via* FLIM z*-*stack imaging at 11.5 h (**Figure 4d**).

## CONCLUSIONS

Monitoring the maturation of condensates to aggregates within a single experimental framework has remained a major challenge in the field. Here, we report a series of viscosity-sensitive fluorescent MRs that can distinguish between the dilute phase, liquid-like condensates and solid aggregates of α-syn, enabling their unambiguous detection and simultaneous imaging through fluorescence lifetime-based measurements. This dual capability allows continuous tracking of the progression from early PS through aggregate formation under these conditions. These findings position the new series of MRs as versatile, rationally designable tools for dissecting the molecular events underpinning protein aggregation in complex biological systems.

## MATERIAL AND METHODS

### Synthesis of sulfo-Cy3-N_3_ derivatives

Unless otherwise stated, all reagents were purchased from commercial suppliers in 99% purity. Fresh stock solutions of 10 mM sulfo-Cy3-N_3_ (**1**), 10 mM CuSO_4_, 1 mM tris(3-hydroxypropyltriazolylmethyl)amine (THPTA), and 10 mM sodium ascorbate, were prepared using HPLC-grade H_2_O. 10 mM stock solutions of the corresponding alkyne, namely 3-ethynylpyridine (**2**), 4-ethynylphenyl acetate (**3**), 3-methoxyphenylacetylene (**4**), 4-methoxyphenylacetylene (**5**), phenylacetylene (**6**), 1-ethynyl-3-(trifluoromethyl)benzene (**7**), and 4-ethynylaniline (**8**), were prepared in DMSO. Alkynes **2– 8** were used to prepare **9–15**, respectively, using the following protocol: 10 mM sulfo-Cy3-N_3_ (1 eq, 10 μL), 10 mM alkyne (10 eq, 100 μL), 10 mM CuSO_4_ (1 eq, 10 μL), 10 mM sodium ascorbate (5 eq, 50 μL), and 1 mM THPTA (5 eq, 500 μL), were pipetted into a 1.5 mL glass vial. The reaction mixtures were made up to 1 mL by using HPLC-grade H_2_O (330 μL). The vials were sealed and placed vertically on a shaker for 24 h at room temperature. The formation of the product was confirmed by liquid chromatography-mass spectrometry, LCMS (see Supporting Information for LCMS traces). Sulfo-Cy3-3-pyridine (**9**) – 96%: MS (ES+) m/z 846.20 ([M+2Na]^+^), 824.10 ([M+Na]^+^), 802.20 ([M]^+^), 401.70 ([M]^2+^); sulfo-Cy3-4-Ph-acetate (**10**) – 95%: MS (ES+) m/z 881.10 ([M+Na]^+^), 859.20 ([M]^+^), 430.20 ([M]^2+^); sulfo-Cy3-3-MeOPh (**11**) – 95%: MS (ES+) m/z 831.20 ([M]^+^), 416.20 ([M]^2+^); sulfo-Cy3-4-MeOPh (**12**) – 95%: MS (ES+) m/z 853.20 ([M+Na]^+^), 831.20 ([M]^+^), 416.20 ([M]^2+^); sulfo-Cy3-Ph (**13**) – 95%:MS (ES+) m/z 823.20 ([M+Na]^+^), 801.20 ([M]^+^), 401.20 ([M]^2+^); sulfo-Cy3-3-CF_3_-benzene (**14**) – 97%: MS (ES+) m/z 891.20 ([M+Na]^+^), 869.20 ([M]^+^), 435.20 ([M]^2+^); sulfo-Cy3-4-aniline (**15**) – 96%: MS (ES+) m/z 838.20 ([M+Na]^+^), 816.20 ([M]^+^), 408.70 ([M]^2+^).

LCMS data was obtained using an Agilent 1260 Infinity II HPLC, equipped with a UV-VIS diode array detector, and G6125B MSD. An Agilent Poroshell HPH-C18 3.0×50mm 2.7μm column was used. Absorption spectra were recorded from 190**–**1100 nm on an Agilent 8453 UV-Vis spectrophotometer. Fluorescence spectra were measured using a Fluoromax-4 spectrofluorometer (Jobin-Yvon, Horiba) and corrected for the detector sensitivity. All samples were excited at 540 nm, and both the excitation and emission slit widths were set to 3 nm. The measurements were conducted in quartz cuvettes with a 1 cm path length.

### Viscosity, polarity, and temperature calibrations

The fluorescence decays of **1**, and derivatives **9–15**, were measured using time-correlated single photon counting (TCSPC) on a Jobin Yvon IBH data station (500 F, HORIBA Scientific Ltd) with a 467 nm NanoLED as an excitation source (FWHM 200 ps) and fitted with either a bi-exponential or tri-exponential function, including a constant offset, using DataStation v2.2 software. An ND30B, Unmounted Reflective ø25 mm ND Filter (optical Density: 3.0) was used for the prompt and an FGL530 - ø25 mm OG530 Colored Glass Filter (530 nm Longpass) was used for test samples. The instrument response function, which was required to fit the decays, was measured by recording a scattering signal from a cuvette with a dilute Ludox® solution. The concentrations of **1** and **9–15** used in all absorption, steady-state and time-resolved fluorescence spectroscopic measurements was 0.2 μM, which gives absorbance of ∼ 0.03 at 548 nm.

To probe the effect of varying viscosity and polarity on the fluorescence lifetimes of **1** and the derivatives (**9–15**) we synthesized, TCSPC was conducted in solutions of varying water:glycerol or water-dioxane content, respectively. Note that for the viscosity calibrations, the percentages of glycerol have been rounded to the nearest decimal place, and that the 100% glycerol sample contains 4 uL of dye dissolved in water. A customized cuvette holder also allowed the sample to be heated to a specific temperature, offering an alternative method of changing the solvent viscosity and a way to determine whether the lifetimes of our dyes also exhibited a temperature dependence. The dynamic viscosities of the water-glycerol mixtures of varying composition were calculated at a given temperature using a freely available online calculator (31).

### α-syn Expression and Purification

Expression and purification of α-syn WT was performed based on an existing protocol (34). To summarize, BL21-Gold (DE3) competent *Escherichia coli* (*E. coli*) cells (Agilent Technologies) were transformed with the pT7-7 α-syn FL plasmid (a gift from Hilal Lashuel, Addgene, USA) (35). Expression of α-syn was induced overnight using 1 mM IPTG at 28 °C. Following centrifugation, the cells were resuspended in buffer A (20 mM Tris-HCl, 1 mM EDTA, EDTA-free Protease Inhibitor Cocktail (Roche, Basel, Switzerland), pH 8.0). The cells were then lysed by sonication and centrifuged. The supernatant was then boiled before another round of centrifugation. Next, the supernatant was treated with streptomycin sulfate (10 mg/mL), followed by further centrifugation. α-syn in the supernatant was precipitated using ammonium sulfate (360 mg/mL) and isolated using a final centrifugation. The pellet was re-suspended in buffer A and further purified by anion exchange chromatography (HiPrep Q HP 16/10 column) using a gradient elution of 1 M NaCl. Finally, the sample was passed through a gel filtration column (HiLoad 16/600 Superdex 75 pg). Pure protein fractions were combined and the concentration measured by UV-Vis spectroscopy, using the absorbance at 275 nm (ε_275nm_ = 5600 M^-1^cm^-1^).

### Condensate and aggregate formation

α-syn PS was induced using a previously described method (24). Briefly, 0 or 60 µM α-syn was incubated with 0 or 25 µM PLK (Sigma-Aldrich) in PS buffer (10% w/v PEG-8000, 0.02% NaN_3_ in PBS, pH 7.4) at 37 °C. The PLK concentration was estimated using an approximate molecular weight of 22, 500 g mol^-1^ (24). Where stated one of the following probes was included in the sample; ThT (10 µM), sulfo-Cy3-N_3_ (**1**) and its derivatives **9–15** (3 µM), DilC_2_(3) (1 µM) or DiSC_2_(3) (3 µM). Samples (100 µL) were loaded into a 96-well glass-bottom plate (Sensoplate, Greiner Bio-One, Austria), sealed with aluminum film to prevent evaporation, and incubated under quiescent conditions for ∼ 60 h in a CLARIOstar Plus microplate reader (BMG Labtech, Germany). At selected time points, incubations were paused to collect images using differential interference contrast (DIC), fluorescence intensity, confocal, or FLIM microscopy. The data were plotted using GraphPad Prism version 10.4.2.

As **9–14** were used without further purification, all contained 1 eq. of CuSO_4_, 5 eq. of Na-ascorbate and 5 eq. of THPTA, and a maximum concentration of 10 eq. of alkynes **2–7** remaining from the click reaction. When dissolved in water in the presence of these reagents, the lifetime of **1** did not vary significantly in the presence or absence of each alkyne (**Figure S11a**), ruling out specific interactions. A slight decrease in the lifetime of **1** was observed in both condensates and aggregates (**Figure S11b**) when we monitored α-syn PS in the presence of **1** and 1 eq. of click reagents (1 eq. CuSO_4_, 5 eq. sodium ascorbate, 5 eq. THPTA).

### Differential interference contrast and fluorescence intensity microscopy

At selected time points the samples incubated in the microplate reader were imaged using a widefield Nikon ECLIPSE Ti2-E microscope (Nikon, Japan). The aluminum film was replaced with a clear, microscopy compatible self-adhesive film (ibiSeal, Ibidi, Germany) and DIC and fluorescence intensity images were acquired using a 60x oil objective and excitation 405 nm and emission 515/30 nm (ThT) or excitation 550 nm and emission 641/75 nm (**1** and **9–15**). In all cases, images were acquired at the bottom of the imaging well.

To quantify changes in fluorescence intensity of each MR within different assemblies, the fluorescence images acquired underwent processing using GA3 software integrated into the NIS-Elements software (Nikon, Japan). Briefly, images were first clarified to remove out-of-focus artifacts and then de-noised. Otsu’s thresholding was then applied to separate the image into foreground (*i*.*e*., condensates or aggregates) and background. The number of pixels and intensity of each pixel was summed for the foreground layer. Subsequently, the mean fluorescence intensity per pixel of the foreground in each image was calculated. This quantifies the mean fluorescence intensity of the probe in the presence of the given species in the sample at that specific time point. The data were plotted using GraphPad Prism version 10.4.2.

### Confocal Microscopy and FLIM Methods

Protein samples were prepared for confocal microscopy and FLIM measurements as described above. We also quantified the lifetime of ThT and **1** in PS buffer including 25 µM PLK, in the absence of protein, for comparison with the lifetimes observed in condensates and aggregates. A Leica TSC SP5 II inverted confocal microscope (Leica Microsystems GmbH, Germany) with a Becker&Hickl FLIM module was used to obtain all FLIM images. FLIM images (256 × 256 pixels) were obtained using a Ti:sapphire pulsed laser source (680–1080 nm, 80 MHz, 140 fs, Chameleon Vision II, Coherent Inc., Germany) synchronized with an internal FLIM detector PMH-100 (Becker&Hickl, Germany). Single photon counting was provided by a TCSPC SPC830 single photon counting card (Becker & Hickl GmbH). All samples containing **1, 9–15**, DiSC_2_(3) or DilC_2_(3) were excited using two-photon pulsed excitation at 960 nm. Emission was collected over 525–700 nm. When ThT was employed as an MR, samples were excited at 880 nm, and emission collected over 460–25 nm. For all samples, the pinhole diameter was set to maximum to achieve optimum light collection efficiency. In all experiments a 20x air-objective (0.7 NA) was used (Leica Microsystem Ltd, Germany). Instrument response functions at a given excitation wavelength were measured using the reflection of the excitation beam from crystals of urea, grown on the glass cover slide. We mitigated any potential temperature effects on our lifetime data by allowing all PS samples to cool to room temperature for 10 minutes before imaging. Apart from the z-stack data presented in **Figure 4d**, FLIM images were acquired at the bottom of the imaging well

### FLIM Data Analysis

FLIM data were acquired for ∼ 1-2 min to achieve a peak photon count of at least 100 for each pixel, after applying a 3 × 3 binning procedure (square bin of 1) to the image. Data analysis and fitting of the resulting images was conducted using SPCImage (v 8.9, Becker & Hickl GmbH, Germany). The time-resolved fluorescence decay, *I*(t), at each pixel was fitted to

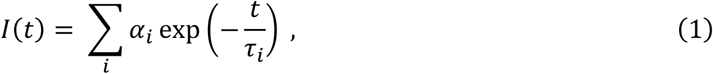

Using Maximum Likelihood Estimation (MLE) algorithm, where α_i_and *τ*_i_ are the pre-exponential factors and lifetimes of the i-th component of the decay, respectively, and t is the time after the laser pulse. The mean fluorescence lifetime, *τ*_m_, was calculated as

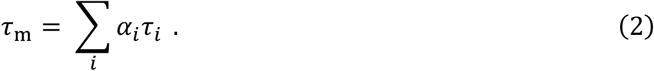

Pixelwise *τ*_m_ distributions over either whole image or regions of interest (condensates, aggreagte), as noted in the relevant figure captions, were then computed from these lifetime values in MATLAB to achieve a quantitative comparison of lifetime changes between different molecular rotors. The parameter, *τ*_m_(max) corresponds to the maximum of the lifetime distribution.

The time-resolved fluorescence decays, over the whole image, were further analyzed by phasor analysis (33) using a home-written MATLAB routine. Briefly, phasor analysis is a model-free method of representing time-resolved fluorescence decays by calculating the real, g(*ω*), and imaginary, *s*(*ω*), parts of the fluorescence decay’s Fourier transform. The real and imaginary components are calculated as,

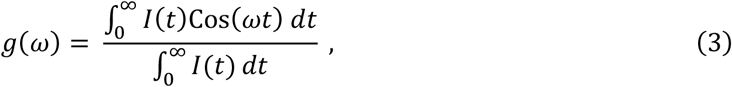

and

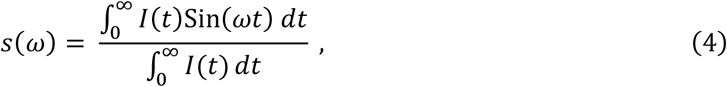

where *ω* is the repetition rate of the exciting laser (80 MHz) in angular frequency units. In this analysis scheme, mono-exponential decays lie on a so-called universal circle, centered at (1/2, 0), with the phasor co-ordinates shifting in a counterclockwise direction on increasing lifetime. Multi-exponential decays lie within the universal circle. If species undergo a sequential conversion (e.g. A → B) then the phasor co-ordinates measured during their interconversion will lie on a straight line.

## ACKNOWLEDGMENTS

All authors gratefully acknowledge funding from the Leverhulme Trust as part of a Research Project grant (RPG-2023-147). N.F. is thankful for a PhD studentship funded as part of the Engineering and Physical Sciences Research Council CDT in Chemical Biology (EP/S023518/1). M.P.P. thanks to the Engineering and Physical Sciences Research Council for a Postdoctoral Open Fellowship (EP/Y021576/1). F.A.A. thanks UK Research and Innovation for a Future Leaders Fellowship (MR/S033947/1) and Fellowship renewal (MR/Y003616/1). We thank Marco Storch and London Biofoundry for providing access to the Nikon Ti2 Eclipse microscope used for fluorescence intensity measurements.

## AUTHOR CONTRIBUTIONS

J.G., N.F., R.J.T., and M.P.P. performed all experiments. All authors conceptualized the work and analyzed the data. F.A.A., R.V., and M.K.K. supervised the study. All authors prepared the manuscript and discussed the interpretations of the results.

