## Supplementary Information for "A Molecular Rotor-Based Platform for Dissecting Protein Phase Separation and Aggregation"

### 1. Comparisons with Commercially Available Molecular Rotors

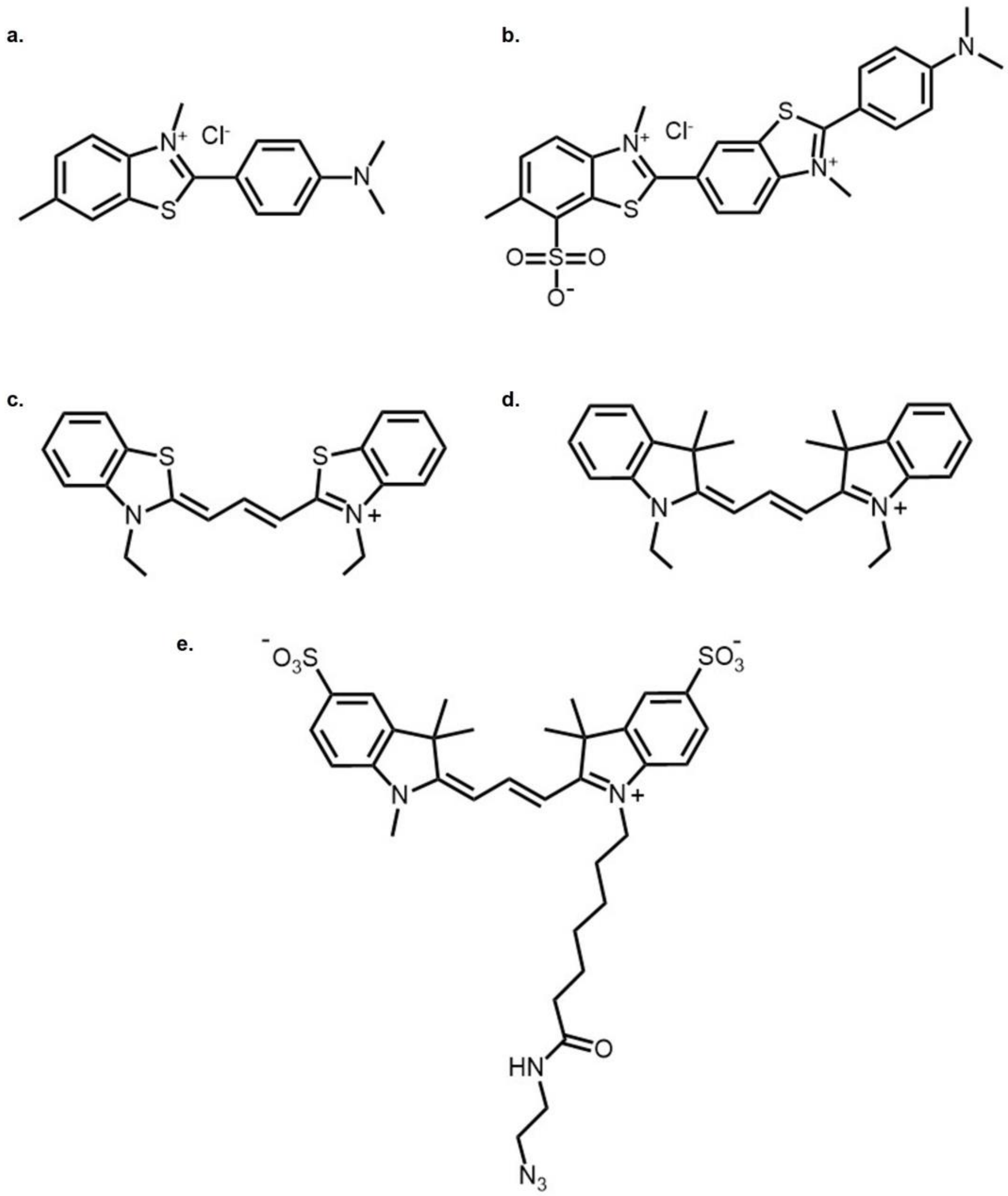

**Figure S1.** Chemical structures of (**a**) ThT, (**b**) ThS, (**c**) DiSC$_2$(3), (**d**) DiIC$_2$(3), and (**e**) Sulfo-Cy3-N$_3$. The structure shown in (**b**) is one component in the mixture of compounds which make up commercial ThS.

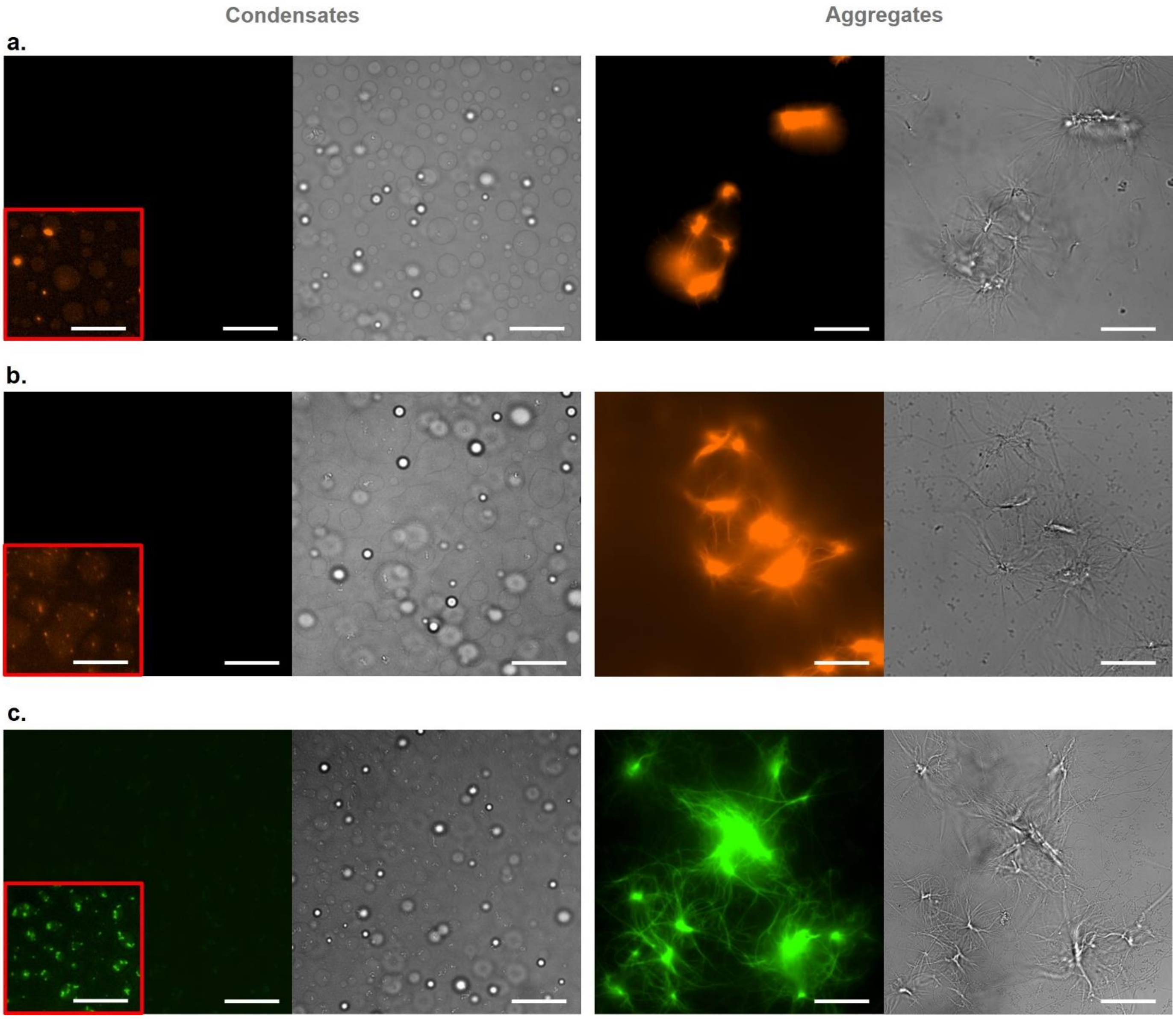

**Figure S2. a-c.** Representative fluorescence intensity (left) and DIC (right) images of condensates (~ 1 h) or aggregates (~ 60 h) formed by the incubation of α-syn in PS buffer, including PLK, in the presence of either (**a**) $DiSC_2(3)$, (**b**) $DiIC_2(3)$, or (**c**) ThS. ThS image acquisition and processing settings are the same as for ThT (**Figure 1b**) while $DiSC_2(3)$ and $DiIC_2(3)$ image acquisition and processing settings match those used for Sulfo-Cy3-$N_3$ (**Figure 1b**). Condensate image inserts have been adjusted to maximum contrast such that any condensate staining would become visible. Scale bars represent 25 μm.

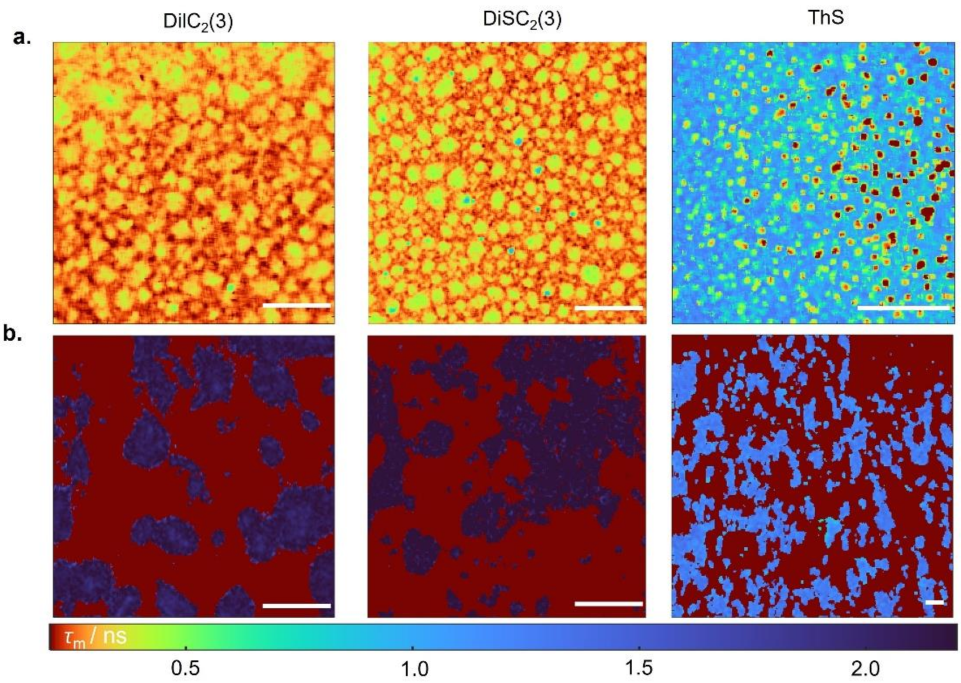

**Figure S3. a,b.** Representative FLIM images of (**a**) condensates (~ 2 h) or (**b**) aggregates (~50 h) formed by the incubation of α-syn in PS buffer, including 25 μM PLK. Samples were incubated in the presence of either $DiSC_2(3)$, $DiIC_2(3)$, or ThS as indicated above the figure. Note the formation of precipitated dye, with a shorter lifetime than in the bulk solution for ThS at 2 h. Scale bars represent 100 μm. The lifetimes observed in condensates can be directly compared with those observed for **13** at 2h in **Figure 4**.

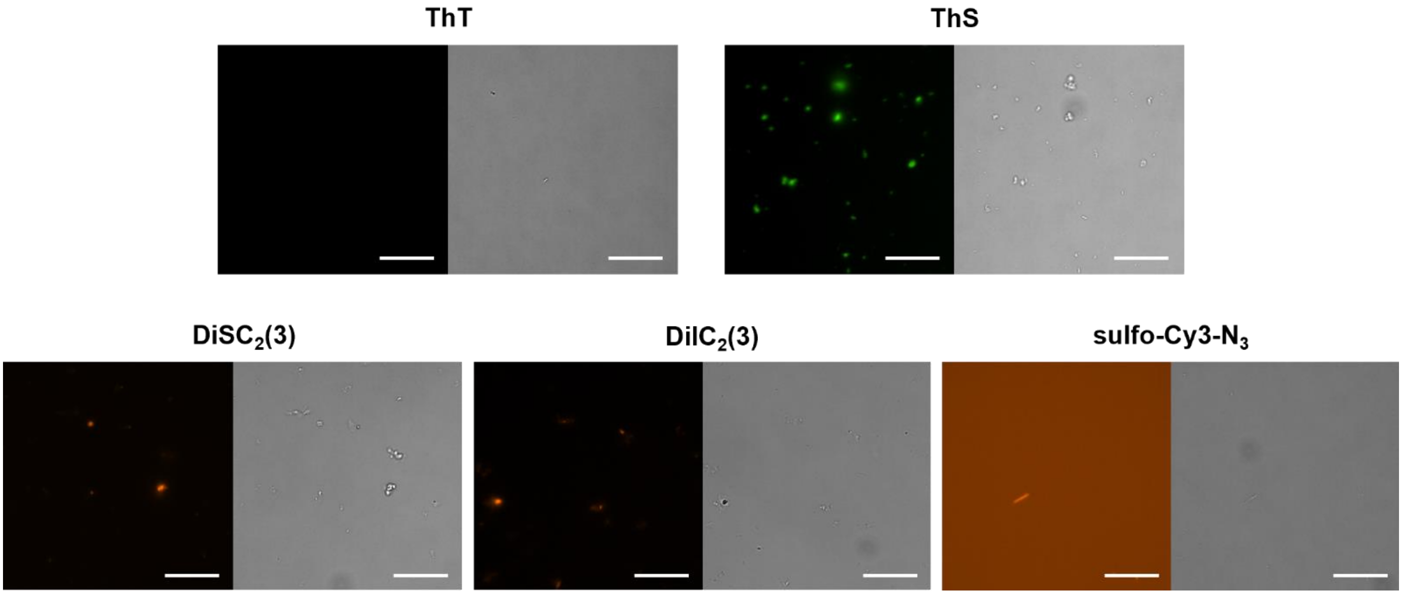

**Figure S4.** Representative fluorescence intensity (left) and DIC (right) images of the commercial MRs incubated in PS buffer alone (i.e. no α-syn or PLK added) for ~ 60 h. ThT and ThS image acquisition and processing settings match those for the ThT PS sample in **Figure 1b** while $DiSC_2(3)$, $DiIC_2(3)$ and sulfo-Cy3-$N_3$ image acquisition and processing settings match those for the sulfo-Cy3-$N_3$ PS sample in **Figure 1b**. Scale bars represent 25 μm.

### 2. Characterization of Molecular Rotor 1 Derivatives

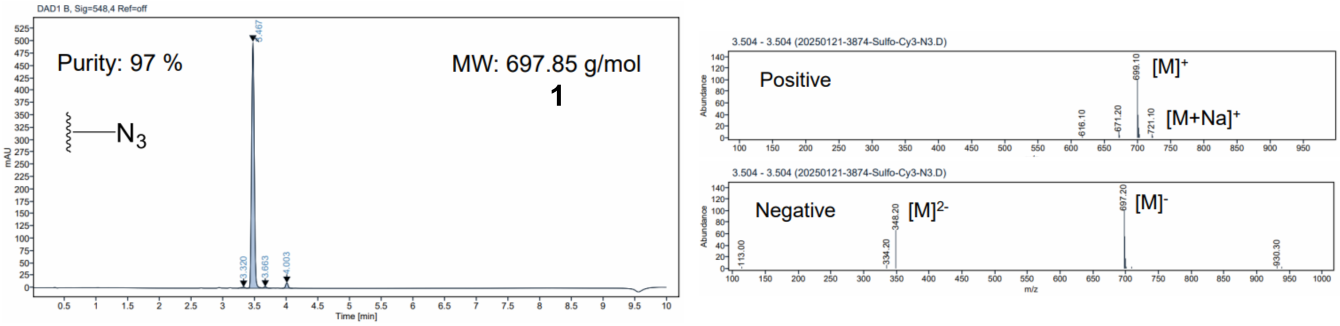

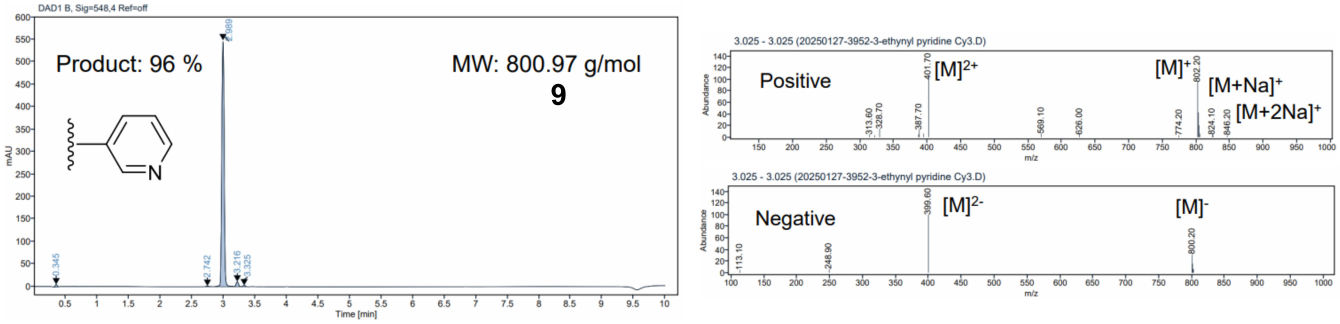

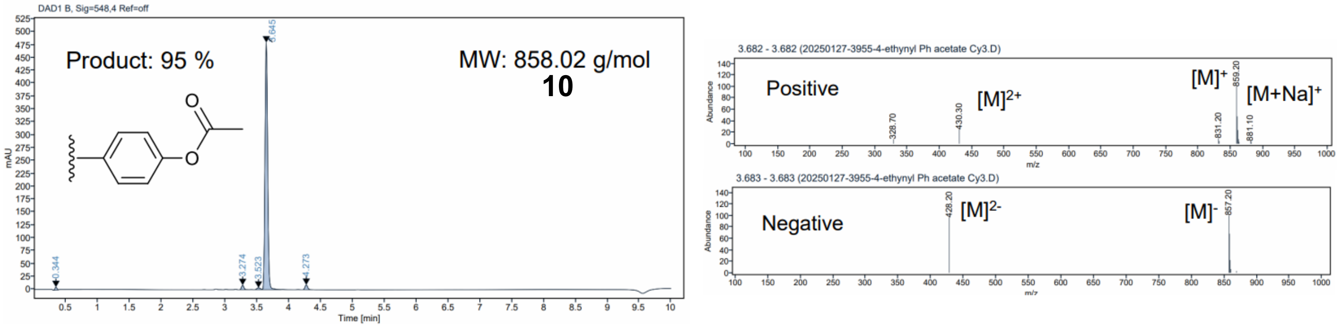

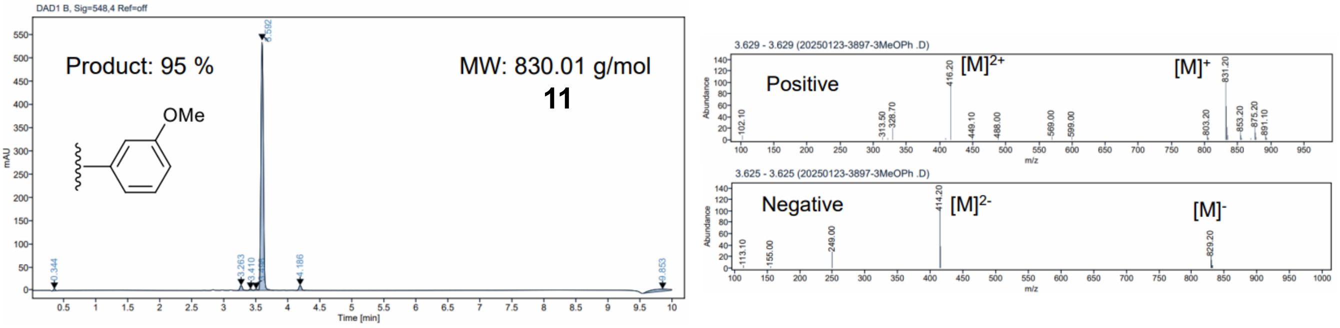

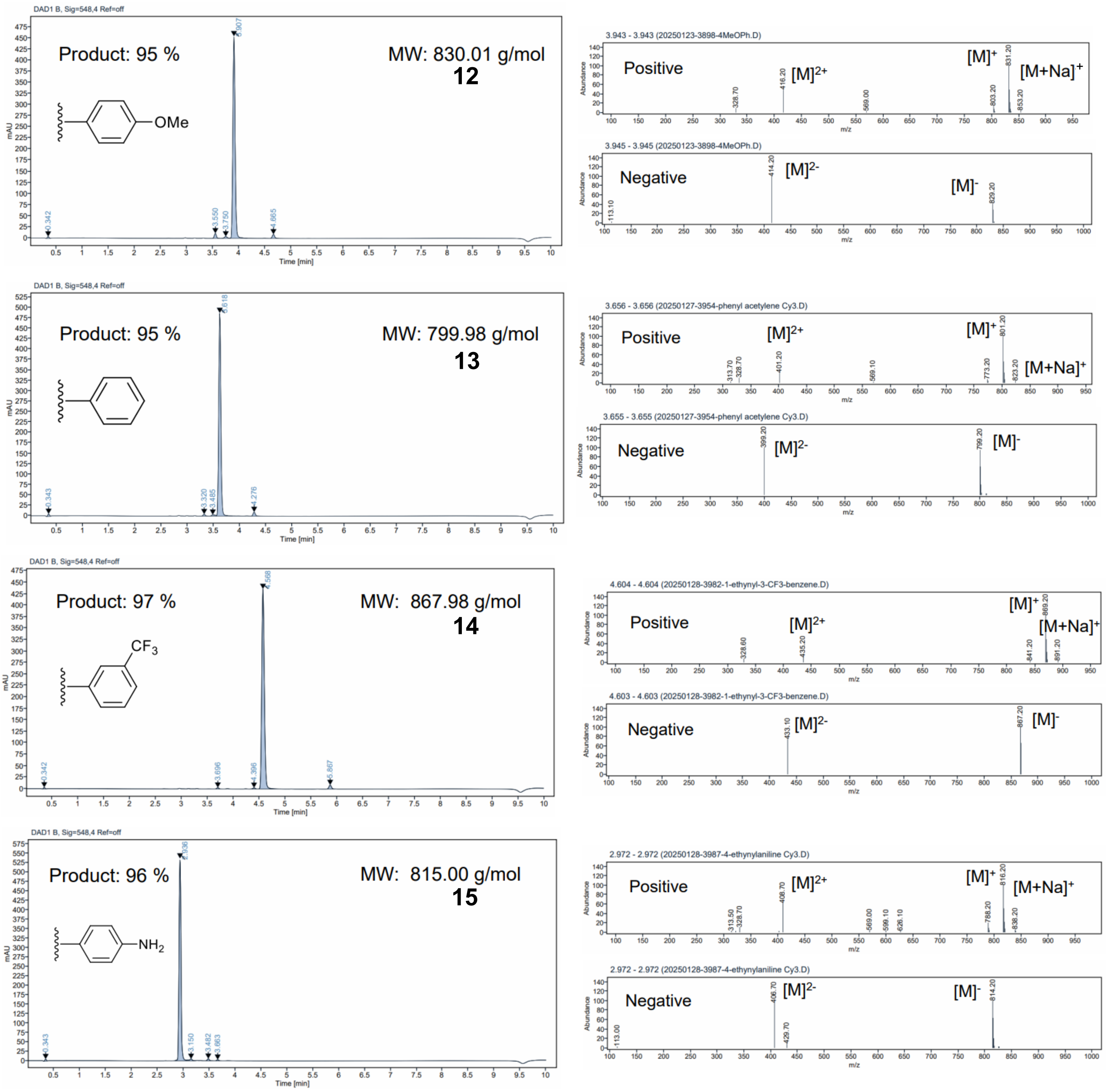

**Figure S5.** UV-vis traces at 548 nm (left) and ES-API data (right) obtained using LCMS analysis of **1** and reaction mixtures of **9–15**. The purity of all derivatives was ≥ 95% immediately after synthesis. As discussed in the main text, the purity of **15** decreased over time and so this compound was deemed unsuitable for further measurements.

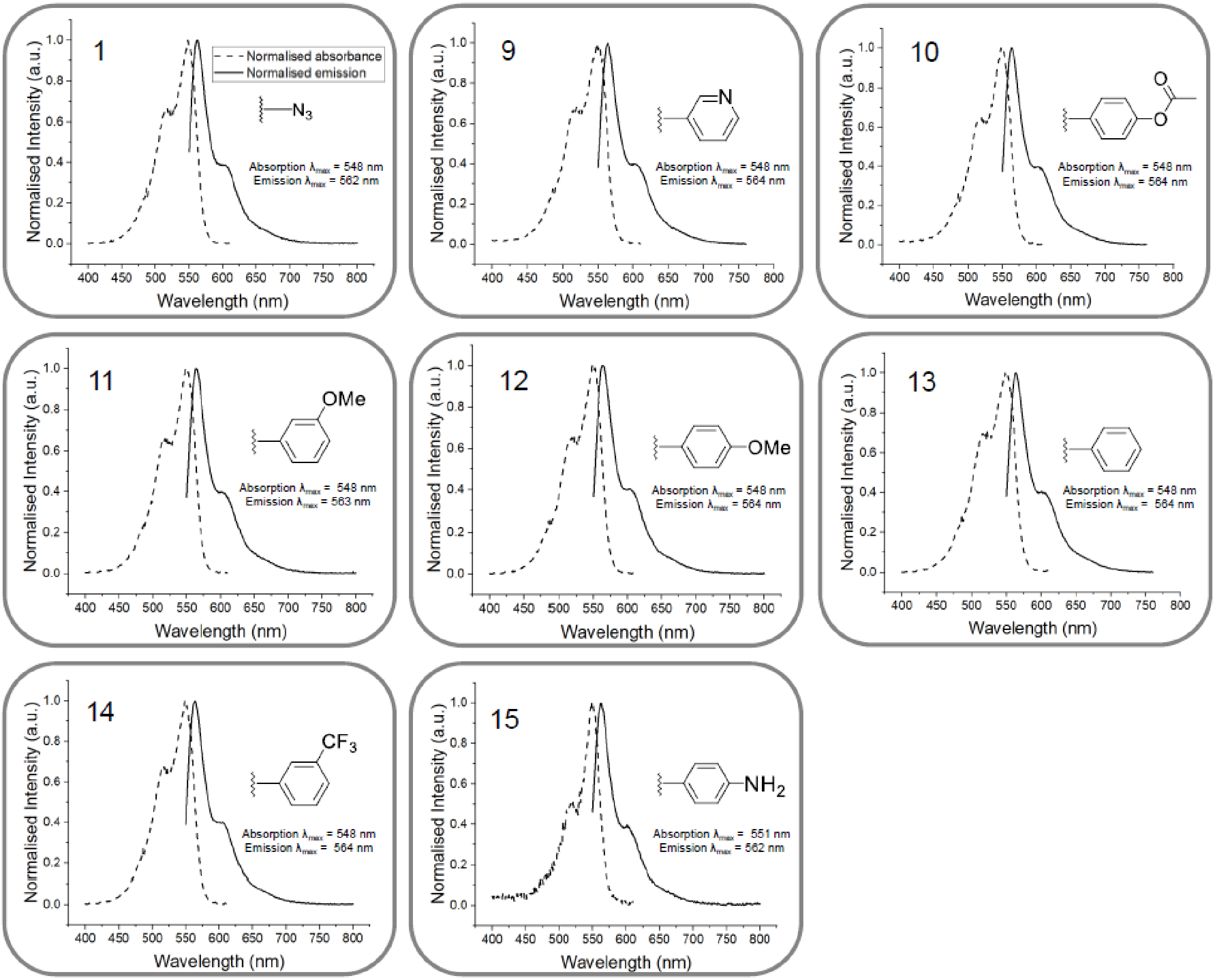

**Figure S6.** The absorption (dashed line) and emission (bold line) spectra of **1** and **9**–**15** (0.2 μM) in PBS. The absorbance and emission spectra maxima are included for each MR.

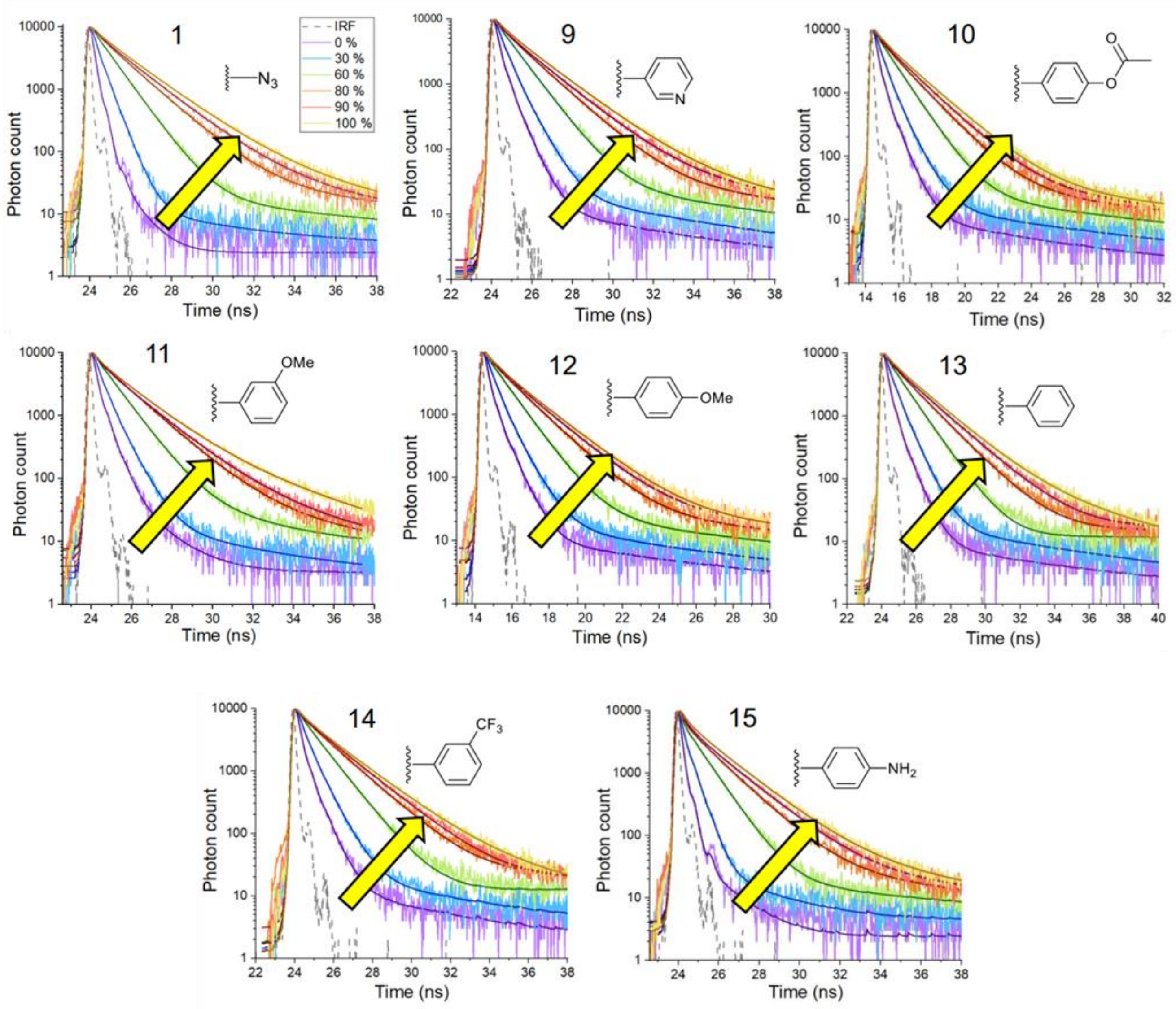

**Figure S7.** Time-resolved fluorescence decays of **1** and **9**–**15** in glycerol/water mixtures of varied ratios (0-100% glycerol, see methods), obtained at room temperature. Increasing viscosity (yellow arrow) increases the fitted fluorescence lifetime of all MRs (see **Table S1** for the mean lifetimes obtained from tri-exponential fits including a constant offset to these data). The fits are shown by solid lines. For reference, the IRF is shown (dashed grey line).

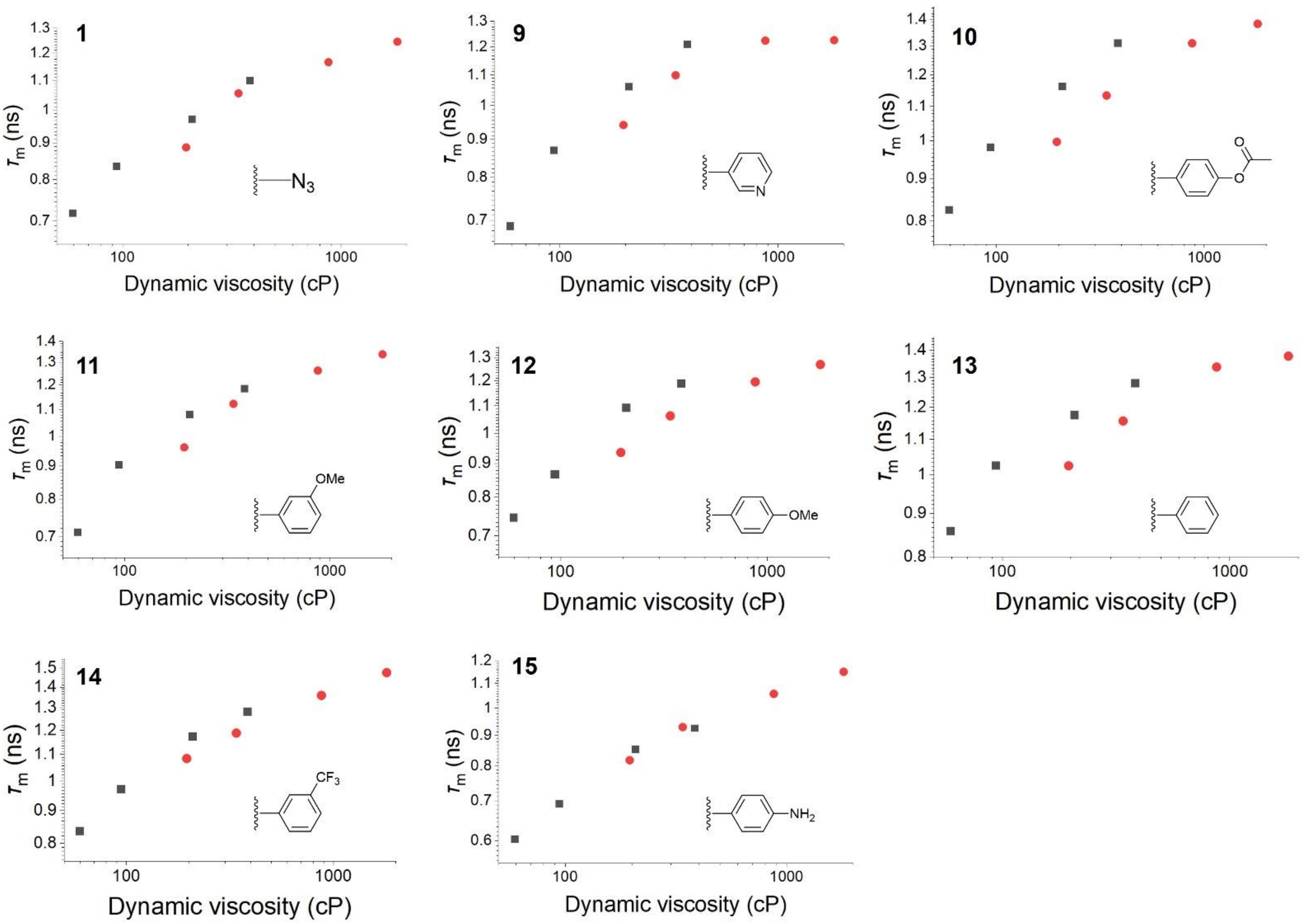

**Figure S8.** Mean fluorescence lifetime ($\tau_\mathrm{m}$) of **1** and **9**–**15** in either a 90% glycerol:water mixture (red) or 100% glycerol (black) measured at 17, 25, 37, and 45 °C. 100% glycerol requires higher temperatures relative to the 90% glycerol:water mixture to achieve the same viscosity. Thus, higher temperatures slightly reduce the fluorescence lifetime of rotors **1** and **9–14** at identical viscosity. The temperature sensitivity of a MR can thereby be inferred from the overlap of the two lifetime curves. From this limited dataset, the lifetime of **15** appears unaffected by temperature.

**Table S1.** $\tau_m$ values, in ns, obtained by fitting either bi- or tri-exponential decays, including a constant offset, to TCSPC data for **1** and **9–15** in solvents of varying water:glycerol ratios. The choice of fitting function was determined from the $\chi^2$ values and fit residuals. The corresponding data and fits used to calculate these values are given in **Figure S7**. The instrumental errors in the fitted parameters are below 5% of the quoted values.

| Viscosity / cP | **1** | **9** | **10** | **11** | **12** | **13** | **14** | **15** |
|---|---|---|---|---|---|---|---|---|
| 0.9 | 0.14 | 0.22 | 0.26 | 0.23 | 0.26 | 0.25 | 0.24 | 0.11 |
| 2.6 | 0.35 | 0.41 | 0.49 | 0.41 | 0.48 | 0.45 | 0.44 | 0.22 |
| 12.8 | 0.72 | 0.78 | 0.80 | 0.78 | 0.75 | 0.81 | 0.82 | 0.44 |
| 66.7 | 1.06 | 1.07 | 1.03 | 1.07 | 1.06 | 1.03 | 0.95 | 0.71 |
| 208.1 | 1.18 | 1.06 | 1.21 | 1.05 | 1.19 | 1.21 | 1.00 | 0.89 |
| 905.7 | 1.40 | 1.25 | 1.37 | 1.25 | 1.37 | 1.31 | 1.30 | 1.04 |

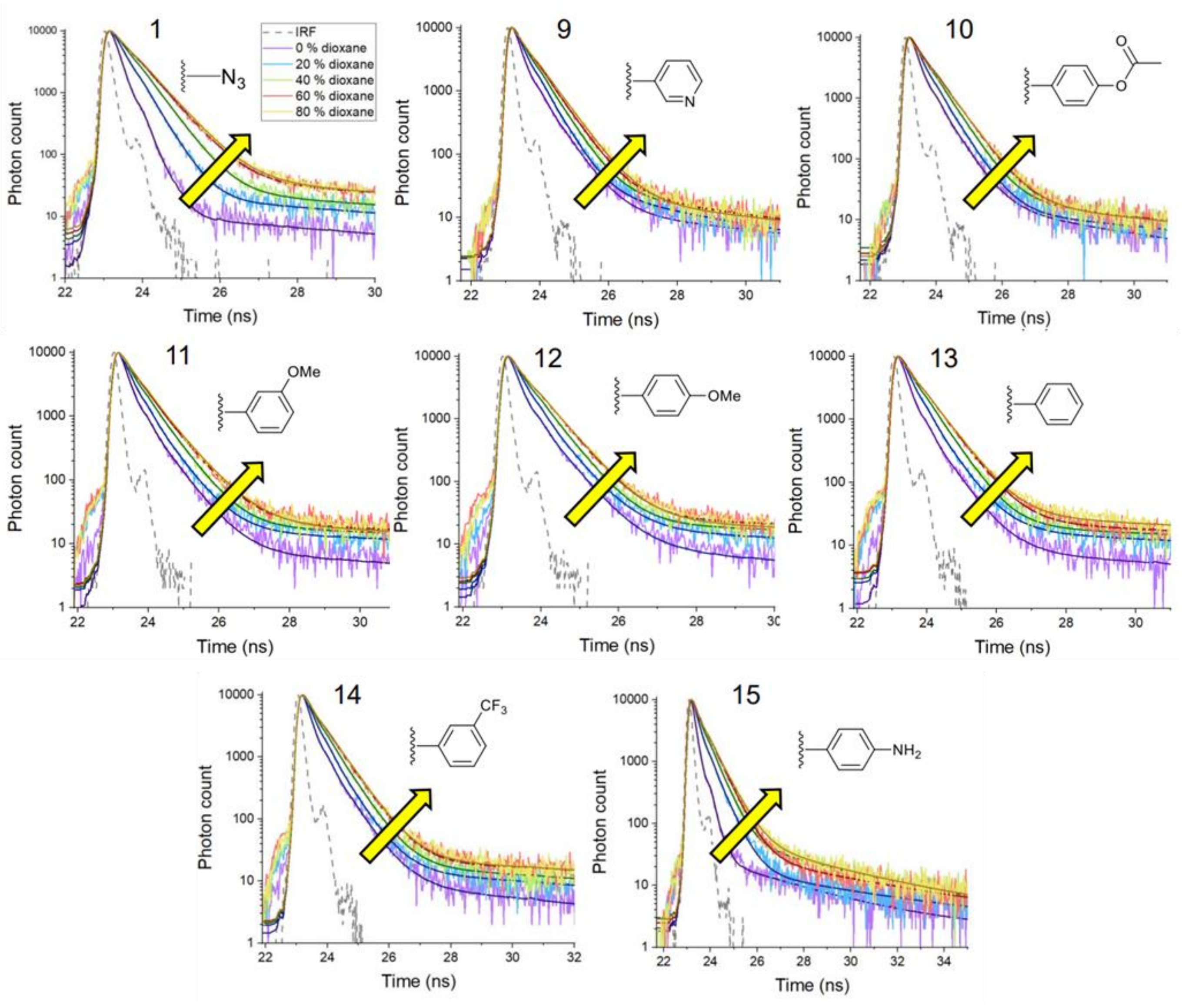

**Figure S9.** Time-resolved fluorescence decays of MRs **1** and **9–15** in varied 1,4-dioxane:water ratios (0-80% 1,4-dioxane) at room temperature. Increased dioxane corresponds to lower polarity. Decreasing polarity (yellow arrow) slightly lengthens the lifetime for all MRs. The fits are shown by solid black lines. For reference, the IRF is shown (dashed grey line).

### 3. Fluorescence Lifetime Distributions for Condensates and Aggregates

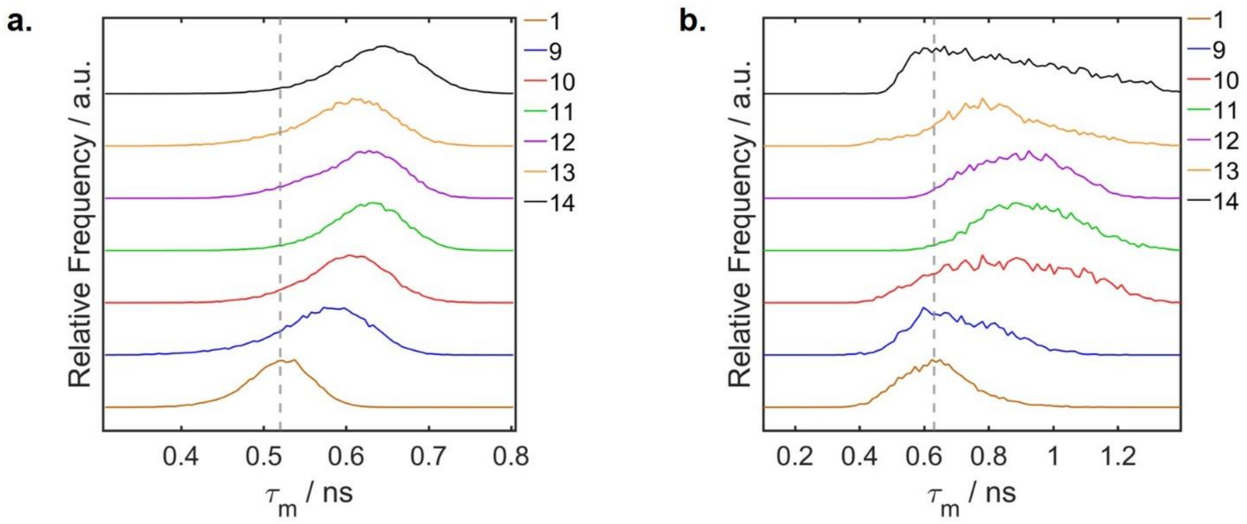

**Figure S10. a,b.** Representative fluorescence lifetime distributions averaged over either (**a**) condensates (~ 6 h) or (**b**) aggregates (~ 60 h) in the images shown in **Figure 3c,** corresponding to PS of α-syn in PS buffer. The maximum of the lifetime distribution corresponding to **1** is indicated by a grey dashed line. Note the increase in lifetimes observed for both condensates and aggregates in the presence of the derivatives **9–14**.

### 4. Controls for Click-Reaction Mixture

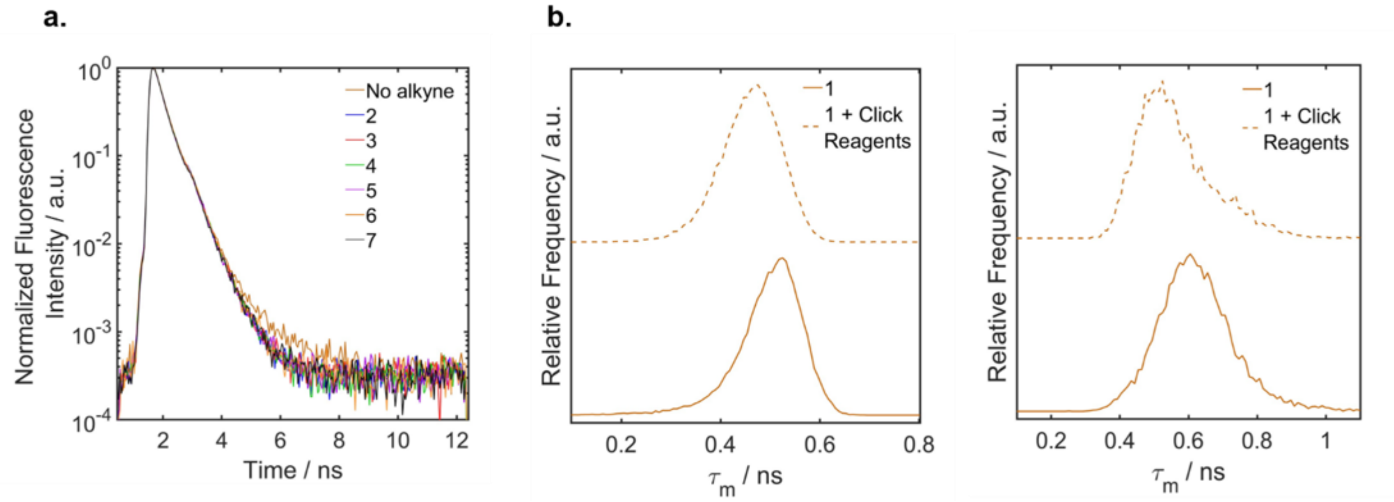

**Figure S11. a**. Time-resolved fluorescence decays obtained following 960 nm excitation, detection over 535-700 nm, for **1** in the presence of 1 eq of click reagents (1 eq $CuSO_4$, 5 eq sodium ascorbate, 5 eq THPTA) dissolved in water. Decays were measured in the absence (orange) or presence of 10 eq of alkynes **2**–**7**, used to synthesize MRs **9–14**. **b.** Fluorescence lifetime distributions obtained for **1** in condensates (left, 6h) and aggregates (right, 50h) of α-syn WT under PS conditions. Lifetimes were recorded in the absence (solid line) or presence (dashed line) of 1 eq of click reagents.
